# Privacy-Aware Kinship Inference in Admixed Populations using Projection on Reference Panels

**DOI:** 10.1101/2022.05.03.490348

**Authors:** Su Wang, Miran Kim, Wentao Li, Xiaoqian Jiang, Han Chen, Arif Harmanci

**Affiliations:** Center for Precision Health, School of Biomedical Informatics, The University of Texas Health Science Center at Houston, Houston, TX 77030, USA; Department of Computer Science and Engineering and Graduate School of Artificial Intelligence, Ulsan National Institute of Science and Technology, Ulsan, 44919, Republic of Korea; Center for Secure Artificial intelligence For hEalthcare (SAFE), School of Biomedical Informatics, University of Texas Health Science Center, Houston, TX, 77030, USA; Human Genetics Center, Department of Epidemiology, Human Genetics and Environmental Sciences, School of Public Health, The University of Texas Health Science Center at Houston, Houston, TX 77030, USA

## Abstract

Estimation of genetic relatedness, or kinship, is used occasionally for recreational purposes and in forensic applications. While numerous methods were developed to estimate kinship, they suffer from high computational requirements and often make an untenable assumption of homogeneous population ancestry of the samples. Moreover, genetic privacy is generally overlooked in the usage of kinship estimation methods. There can be ethical concerns about finding unknown familial relationships in 3^rd^ party databases. Similar ethical concerns may arise while estimating and reporting sensitive population-level statistics such as inbreeding coefficients for the concerns around marginalization and stigmatization. Here, we make use of existing reference panels with a projection-based approach that simplifies kinship estimation in the admixed populations. We use simulated and real datasets to demonstrate the accuracy and efficiency of kinship estimation. We present a secure federated kinship estimation framework and implement a secure kinship estimator using homomorphic encryption-based primitives for computing relatedness between samples in 2 different sites while genotype data is kept confidential.

## Introduction

Genetic relatedness or kinship between two individuals is the probability that two alleles at a random position in the genomes of the individuals are identical-by-descent (IBD), i.e., they are inherited from the same ancestor [1,2]. The kinship coefficient is closely related to other metrics such as the inbreeding coefficient [3] and IBD-sharing probabilities [4], which are essential for estimating population-level genetic information. Kinship estimates are central in behavioral science [5], human evolution [6], linkage mapping studies [7], and association studies [8–10] for the correction of biases caused by cryptic relatedness [9,11]. Numerous computational methods are developed to estimate kinship from marker genotypes but privacy and ethical concerns are sidelined. Kinship statistics are sensitive to individual privacy as they can be used to detect relatives in 3^rd^ party databases without the consent of the owners, for example, by law enforcement [12,13]. Similarly, population-level inbreeding estimates can cause marginalization and stigmatization risks [14–16]. In addition, it is well known that genetic data is very identifying due to its high dimensionality [17–20] and numerous “attacks” have demonstrated that databases can be linked [21–23] to reveal sensitive information. Similarly, genotypes can be recovered [24–26] and sensitive phenotypes can be inferred [27–31] using a small number of marker genotypes. Much of these attacks implicate and create discrimination and stigmatization risks to individuals and their families [32–35]. Therefore genetic kinship estimation presents numerous unaccounted challenges regarding individual and kin privacy [32,34,36].

Kinship estimation methods can be broadly divided into four categories [37]. Moment estimators such as KING [38], REAP [39], plink [40], GCTA [41], GRAF [42], and PC-Relate [43] use identical-by-state (IBS) markers and genotype distances to estimate expected kinship statistics. Maximum-likelihood methods (Such as RelateAdmix [44] and ERSA [45]) use expectation-maximization (EM) to jointly estimate the kinship statistics. Recent methods (such as RAFFI [46], IBDKin [47]) use fast algorithms to search for IBD matches from phased genotypes and estimate kinship from shared IBD estimates. There are also methods that estimate kinship from next-generation sequencing data, which are especially useful from low-coverage sequencing approaches (NGSRemix [48], LASER [49], SEEKIN [50]). While most methods can accurately estimate kinship for individuals with homogeneous ancestry, this is not a tenable assumption in admixed populations[2,51]. Moreover, non-random mating, i.e., assortative mating, among similar ancestral groups [52,53] may bias estimates of kinship. Methods that assume random mating or simple homogeneous populations are not effective in appropriately estimating kinship and may impact downstream analysis and interpretations. Several methods have been proposed for privacy-aware analysis of ancestry and admixture. PREMIX [54] computes admixture rates in a privacy-preserving manner using SGX-based extensions, which are currently deprecated on consumer-side processors. He et al. combined a genome sketching technique with cryptographic evaluation to search for relatives [55]. Similar sketching techniques have been proposed for fingerprint and relative search analysis [56]. Dervishi et al. proposed privacy-aware kinship estimation by integrating local differential privacy and genotypic data hiding [57], which may hinder the utility of genetic data. While these methods are promising, the impact of admixture is not generally taken into account, and the methods are evaluated only for one kinship statistic that provides partial information about relatedness.

Here, we present SIGFRIED, a projection-based approach to utilize existing reference genotype datasets for estimating admixture rates for each individual and use these estimates for kinship and related statistics [49] in admixed populations. The modular formulation of SIGFRIED enables an efficient secure implementation. Usage of component analysis and reference populations with a “distance-based” estimation of admixture has shown promise in previous studies [58,59]. We capitalize on these and propose an efficient approach to estimate kinship, inbreeding, and IBD sharing probabilities. In comparison to previous methods, SIGFRIED imposes less computational burden without the requirement of compute-intensive admixture estimates, which are prohibitively challenging in secure implementations. We implemented a secure federated kinship estimation among 2-sites wherein genetic data is kept confidential while kinship statistics are estimated. Our implementation relies on homomorphic encryption [60], which enables processing encrypted genotype data directly without ever being decrypted and therefore provides provable security guarantees on the genetic data.

## Results

We first describe an overview of SIGFRIED’s estimation approach present accuracy results. After the new kinship estimation results, we formulate the secure implementation and present the secure collaborative kinship analysis scenario.

### Kinship Estimation using Projection-based Admixture Estimates

Figure 1 summarizes the kinship estimation approach by SIGFRIED. Kinship estimation takes a query genotype matrix that contains *S* individuals for which *S* × *S* kinship related statistics are computed. SIGFRIED utilizes principal components and representatives computed from a reference population panel that contains *S*_*tot*_ individuals and *n*_*ref*_ populations. The reference panel genotype is first decomposed into components and for each population, we compute a representative sample. Given the query genotype matrix for *S* individuals, we project the genotypes to the top components of the reference panel computed in the previous step. We next compare each sample to the representatives and assign admixture rates using a non-linear function of genotypes. We finally assign the allele frequencies, *μ*, using the admixture rates. These operations have efficient secure implementations in homomorphic encryption [60] and can be justifiably used in this scenario [61].

**Figure 1.**
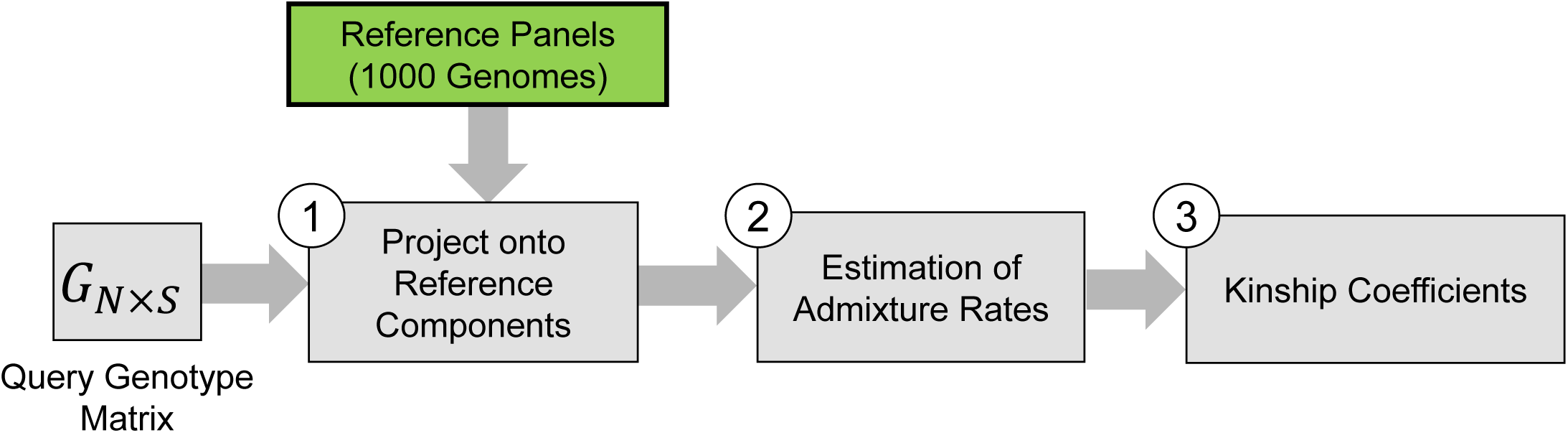
**(a)** Block diagram illustrates the steps for computation of kinship coefficients.

#### Kinship Coefficients

We implement two kinship coefficients. First is the correlation metric, *ϕ*^(corr.)^ *= ρ*(*G*|*μ*), between the individuals. The second kinship metric we use is a novel genotype distance-based metric, *ϕ*^(Dist.)^ = Δ(*G*|*μ*), which integrates individual-specific allele frequencies. For a privacy-aware implementation, the distance and correlation-based can be computed using different strategies. Sites must share the genotypes and allele frequencies. Allele frequencies do not immediately reveal genetic information but they correlate significantly with actual genotypes and may need to be encrypted. These statistics can also be computed in parallel and the final kinship statistic can be aggregated at each site locally.

#### Zero-IBD Sharing Probability

We also report the moment estimator for zero IBD-sharing probability among individuals, which is derived from the expected number of zero identical-by-state (IBS) values:

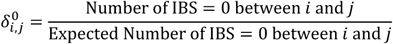

where 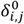 denotes the probability of zero-IBD sharing among individuals *i* and *j*.

#### Inbreeding Coefficient

The inbreeding coefficient for each individual can be estimated from the correlation-based kinship estimator using the established relationship between kinship and inbreeding coefficients:

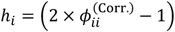

where 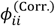 denotes the self-kinship coefficient and *h*_*i*_ denotes the inbreeding coefficient for *i*^*th*^ individual.

### Parameter Selection

We evaluated the impact of the number of variants in the estimation of kinship statistics. For this, we simulated 50 homogeneous pedigrees and computed kinship statistics using SIGFRIED within each pedigree using an increasing number of variants from 500 variants up to 150,000 variants. As the number of variants is increasing, the variance of kinship estimates decreases for each respective degree of relatedness. Figure 2c shows that adding more than 50,000 variants does not provide much change in the variance of the estimated kinship. Qualitatively, as small as 20,000 variants are sufficient for distinguishing 1^st^ and 2^nd^-degree relatives.

**Figure 2.**
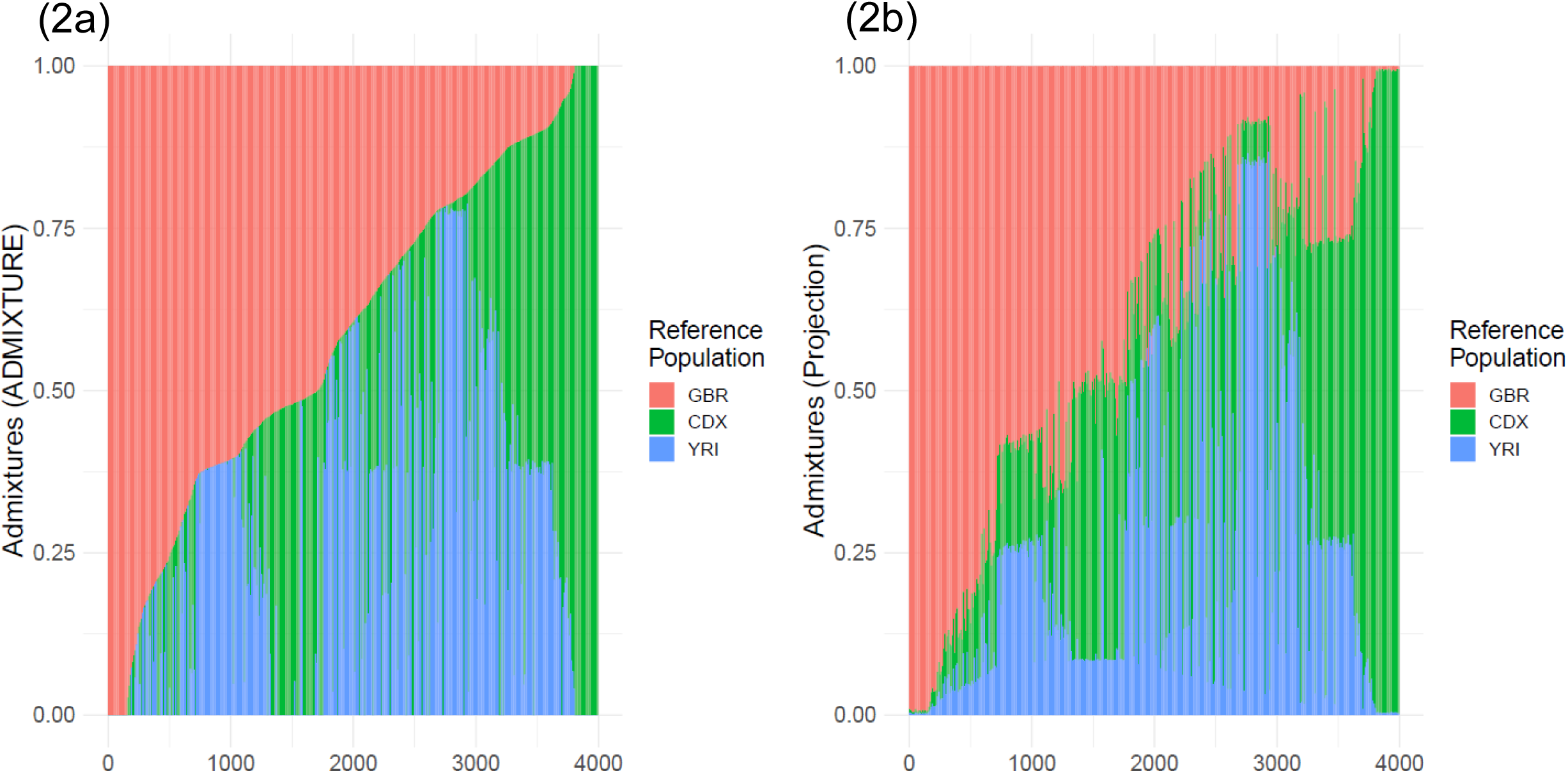
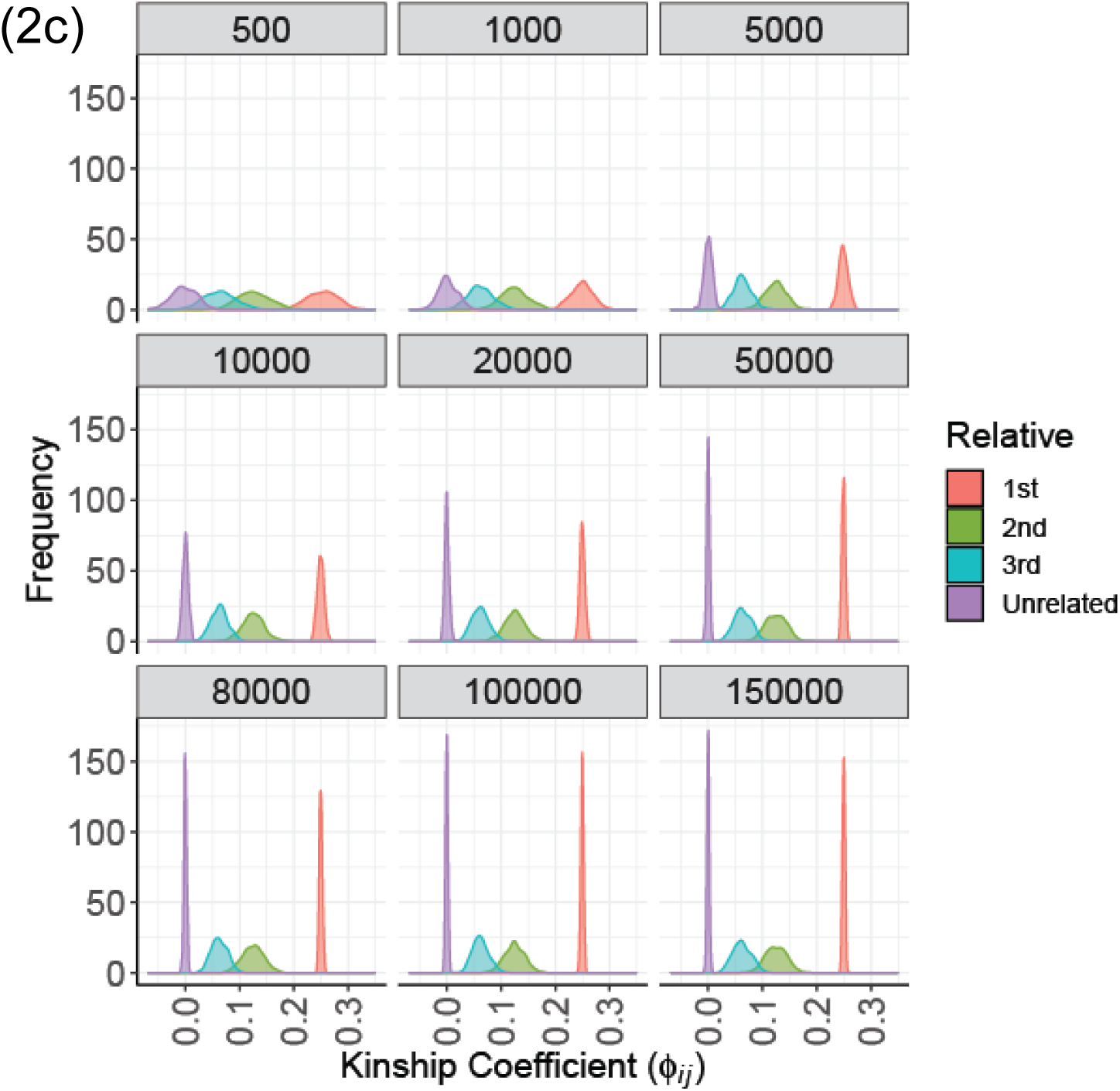
The distribution of assigned admixture rates by ADMIXTURE **(a)** and by projection-based admixture estimation **(b). (c)** The kinship coefficient (x-axis) distribution with different number of variants. Each plot shows a kinship distribution generated using number of variants indicated at the label.

### Comparison of Methods

We compared the correlation and distance-based kinship estimators under homogeneous and heterogeneous pedigree scenarios. We mainly focused on comparing the approaches of SIGFRIED with REAP (with ADMIXTURE tool) and KING-Robust. For SIGFRIED, we use the correlation-based estimator and the projection-based admixture rate estimation to compute individual-specific allele frequencies. We also compare correlation-based and distance-based estimators using allele frequencies estimated by assuming uniform admixture rates over the reference populations, and by using the pooled reference as a single panel to estimate the variant allele frequencies.

### Kinship Estimates in Pedigrees from Same Ancestry

We simulated 500 independent pedigrees where the founding members are randomly selected from a single European population among The 1000 Genomes Project samples. Within each pedigree, we computed the kinship and zero-IBD sharing probabilities between all pairs of members using KING-Robust[38], REAP, and the distance and correlation-based kinship and zero-IBD sharing probability statistics. For SIGFRIED’s projection-based admixture estimates, we used 3 populations from the 1000 Genomes Project as the reference populations to ensure that the admixture estimation step is not trivially applied to a single reference. Overall, we observed that all correlation-based and distance-based methods performed similarly to assign the expected kinship and zero-IBD sharing probability estimates for different levels of kinship (Fig. 3). One observation is that distance-based estimators provide tighter estimates of kinship (Fig. 3c, d), compared to the correlation-based estimators (Fig. 3a, b). Considering that distance-based estimators also have lower computational requirements, these results suggest that they may be more suitable than correlation-based estimators for samples with homogeneous ancestries.

**Figure 3.**
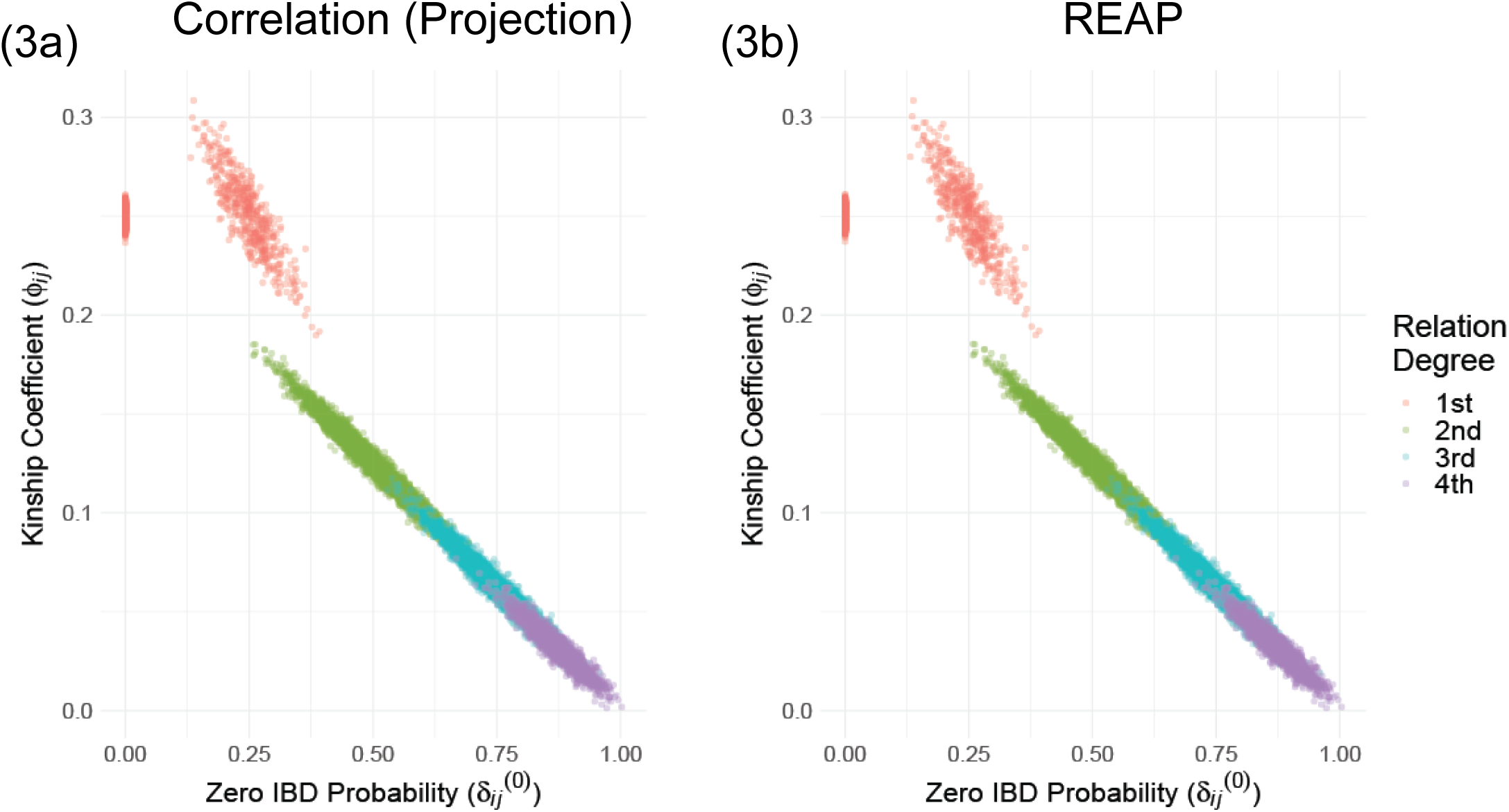
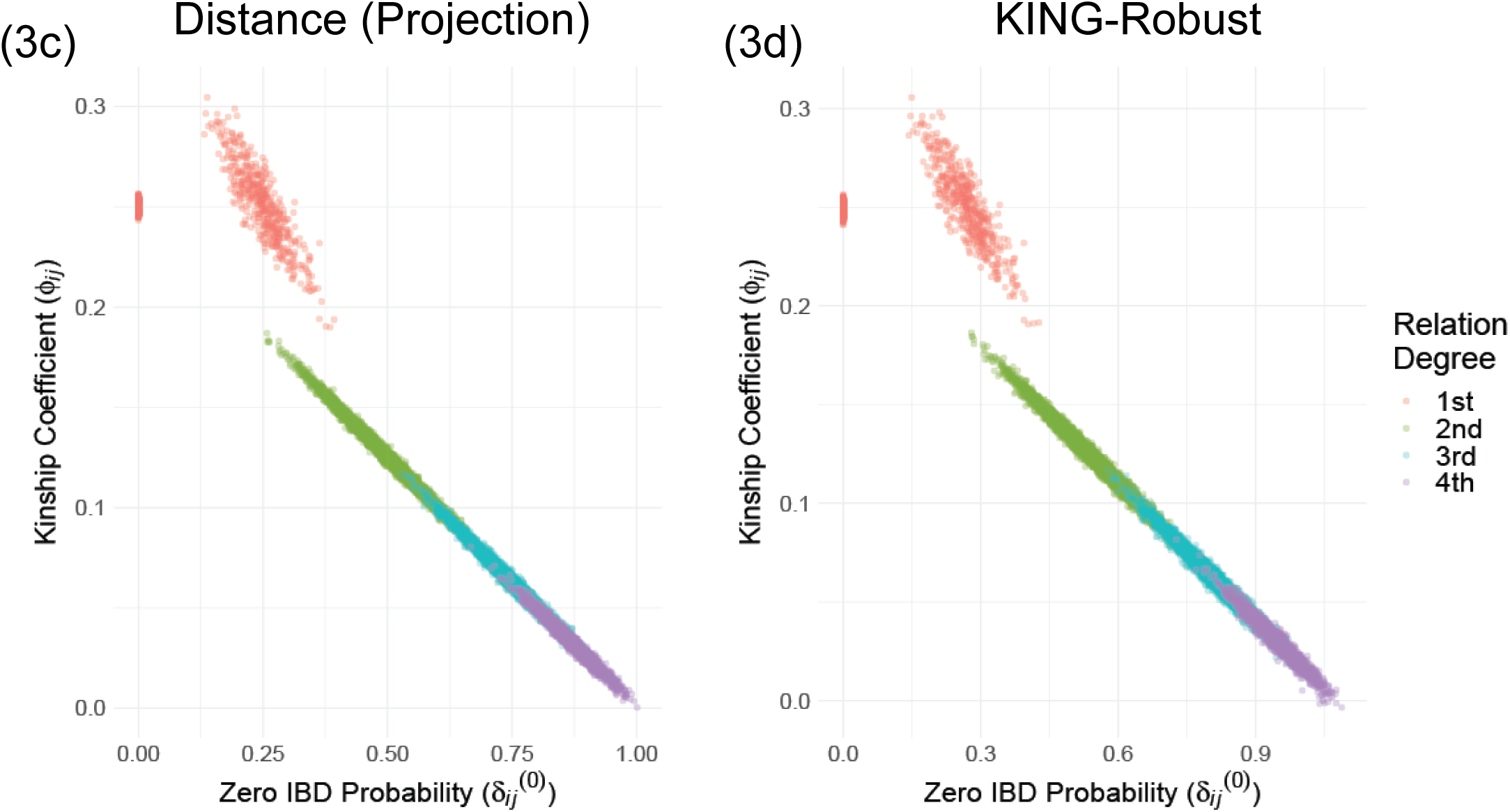
Scatter plots of kinship coefficients in 500 pedigrees from homozygous ancestries. **(a)** kinship coefficient (y-axis) versus Zero-IBD sharing probabilities (x-axis) by SIGFRIED. **(b)** REAP estimates. **(c)** Distance-based estimates. **(d)** KING-Robust estimates.

### Kinship Estimates in Pedigrees from Admixed Ancestry

We next tested the estimation of kinship in admixed ancestries. In the simulation, the founders were selected randomly from populations of European, East Asian, and African descent in The 1000 Genomes Project. For admixed ancestries, we compared the correlation-based estimator using the admixture rates estimated by ADMIXTURE and also with a uniform assignment of admixtures that is equally distributed among 3 reference populations as a control method. We also compared the distance-based estimator with projection-based admixture rates and KING-Robust. In comparison, projection-based estimators and ADMIXTURE-based estimators provide the most accurate results for relatives up to 4^th^ degree (Fig. 4a). KING-Robust underestimates the kinship coefficient, especially for unrelated individuals. Our distance-based estimator largely corrects the negative and heterogeneous trend of KING-Robust. The distribution of kinship coefficients (Fig. 4) indicates that the correlation-based estimators provide single exact peaks around the expected kinship values (Fig. 4b, c). Our novel distance-based estimator exhibits single peaks except for unrelated individuals, for which there is a second peak in negative values. On the other hand, KING-Robust exhibits a fairly high deviation from the expected values with no clear peaks (Fig. 4d, e), which demonstrates the advantage of using a modified distance metric. A similar heterogeneous distribution of kinship is observed for correlation-based estimators that use the pooled reference sample or uniformly assigned admixture rates (Fig. 4f, g).

**Figure 4.**
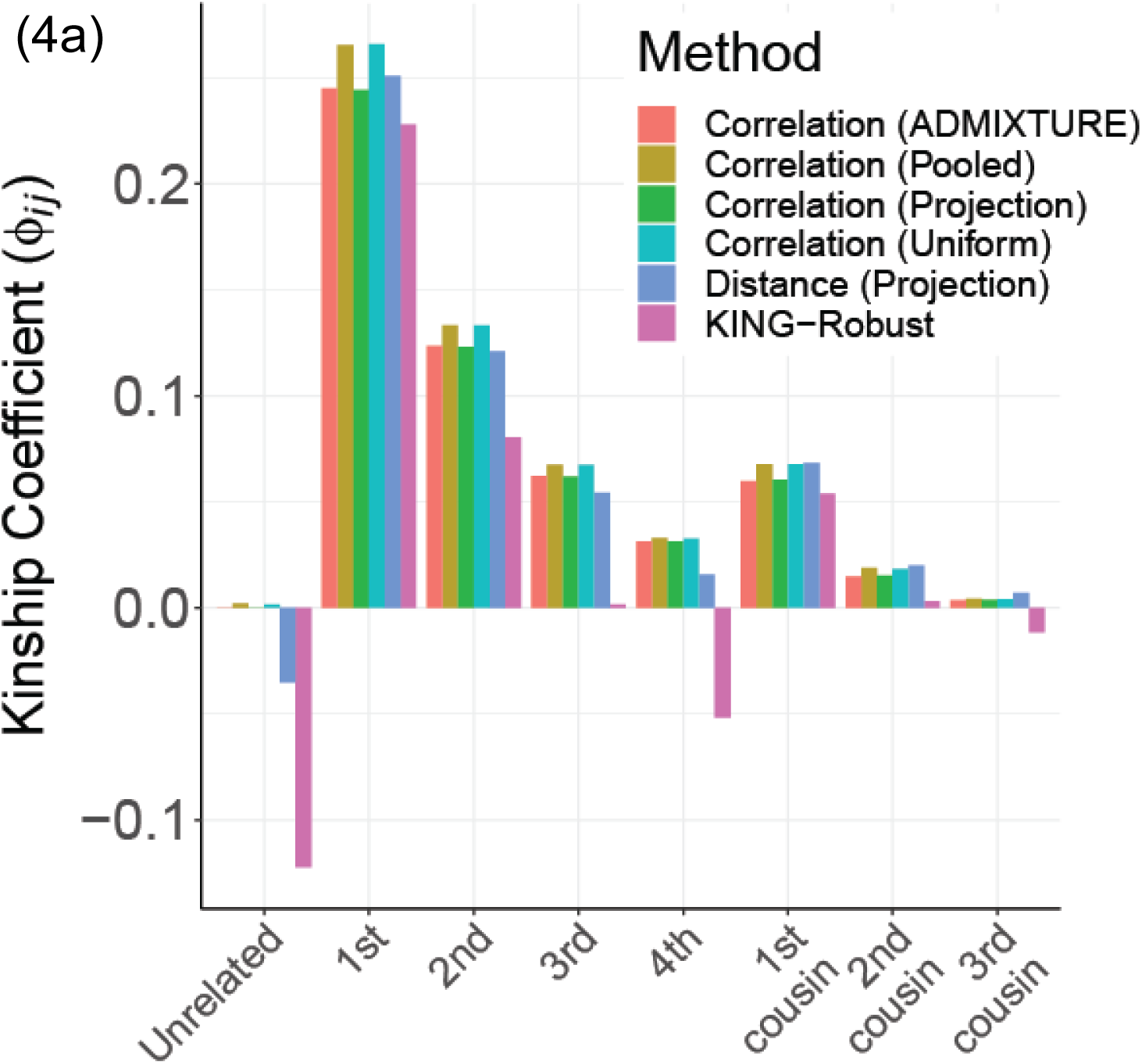
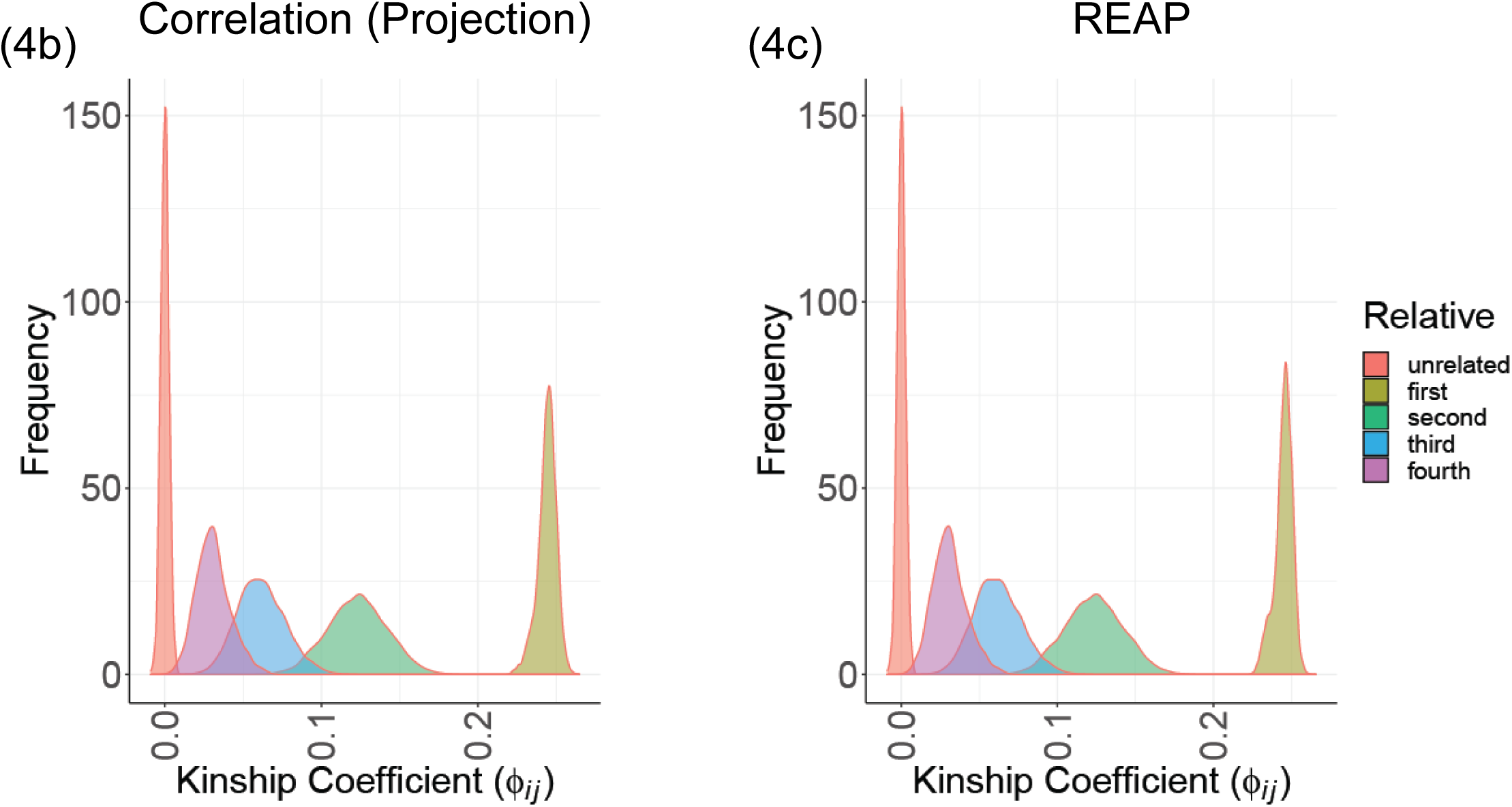
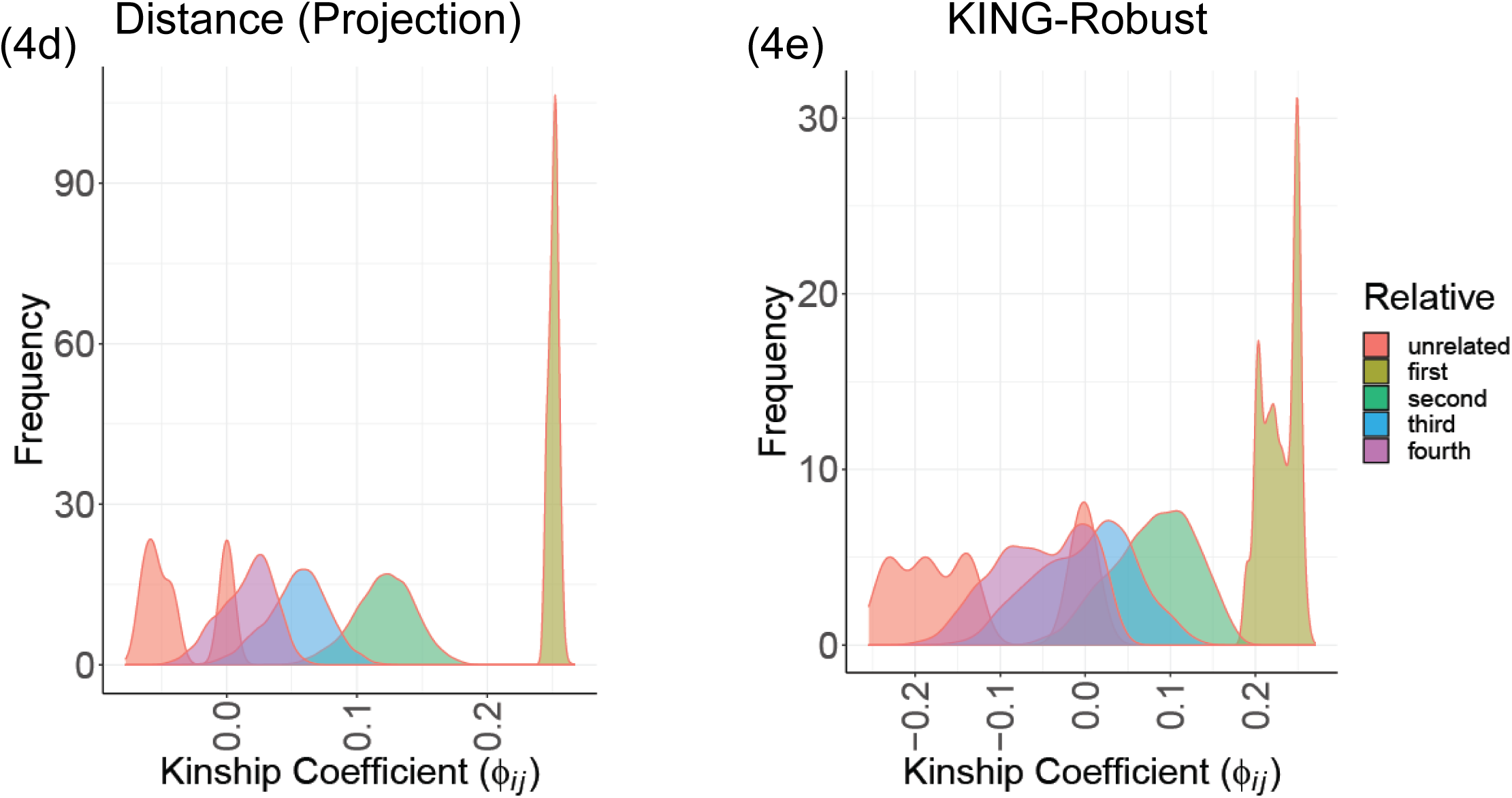
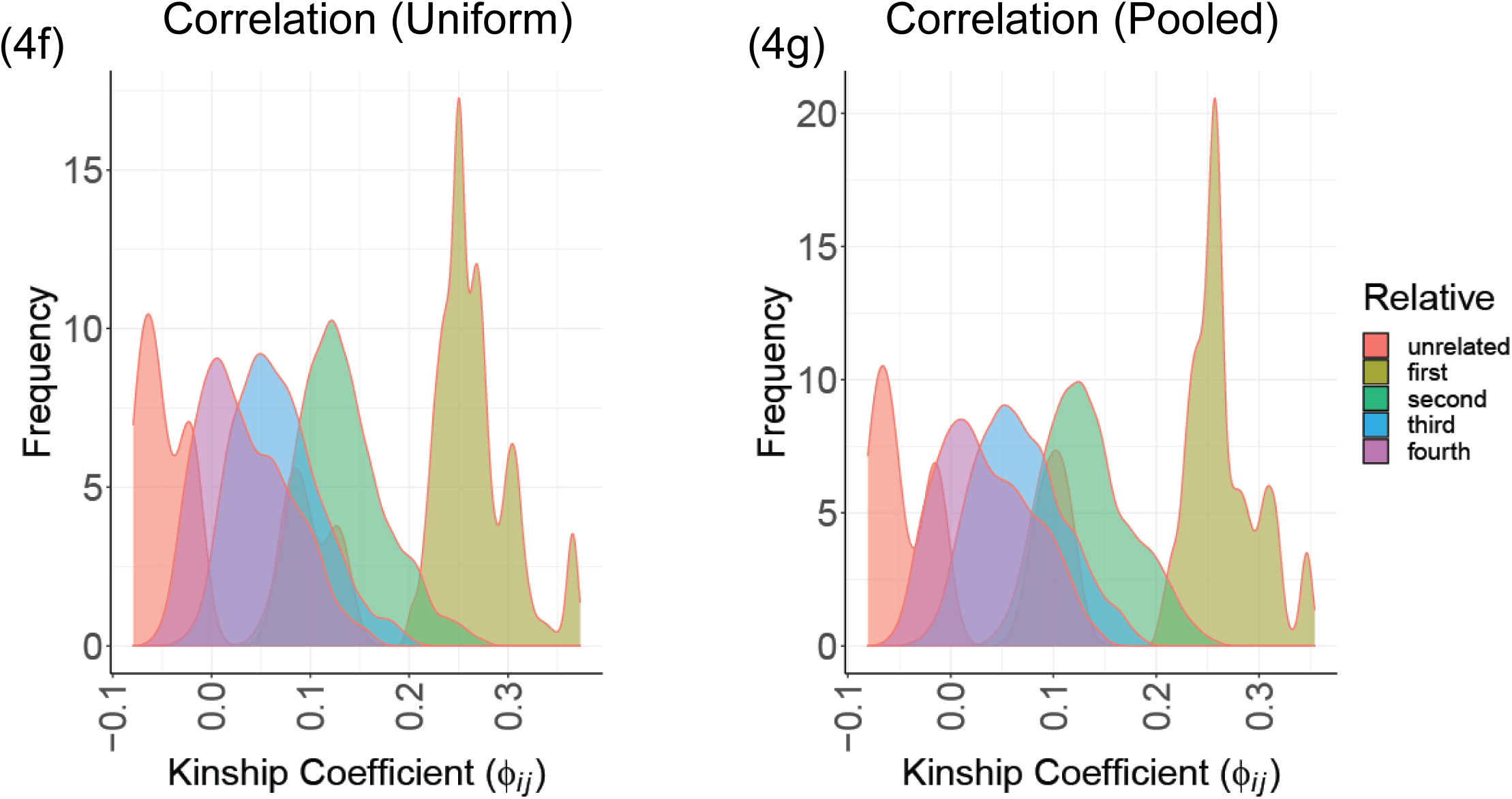
**(a)** Average kinship coefficient estimated by each method for heterogeneous populations. **(b)** Distribution of SIGFRIED kinship estimates. **(c)** Distribution of REAP estimates. **(d)** Distribution of distance-based kinship estimates. **(e)** Distribution of kinship estimates from KING-Robust. **(f**,**g)** Distribution of correlation-based kinship estimates using uniform and all-population admixture assignments for every sample.

### Time and Memory Requirements

We next compared the time and memory requirements of the estimators. To compare the resource requirements of the methods, we estimated the memory and time requirements of SIGFRIED, REAP-ADMIXTURE, and KING-Robust. For all methods, we measured the total time required for admixture estimation, and kinship statistic computations and also the peak memory required for these steps. Overall, KING-Robust runs the fastest and uses the smallest amount of memory (Fig. 5a, b). REAP-ADMIXTURE runs the slowest wherein the majority of time is spent on the estimation of the admixture rates by ADMIXTURE. SIGFRIED runs at least 3 times faster than REAP-ADMIXTURE’s workflow. To test the way that methods scale with the number of reference populations, we compared the resource usage by increasing the number of reference populations (Fig. 5a, b). REAP-ADMIXTURE’s runtime exhibits approximately linear increase in the number of reference populations. On the other hand, SIGFRIED shows a sublinear increase. This indicates that for large admixed populations SIGFRIED’s projection-based approach can provide good accuracy with less computational resource requirements.

**Figure 5.**
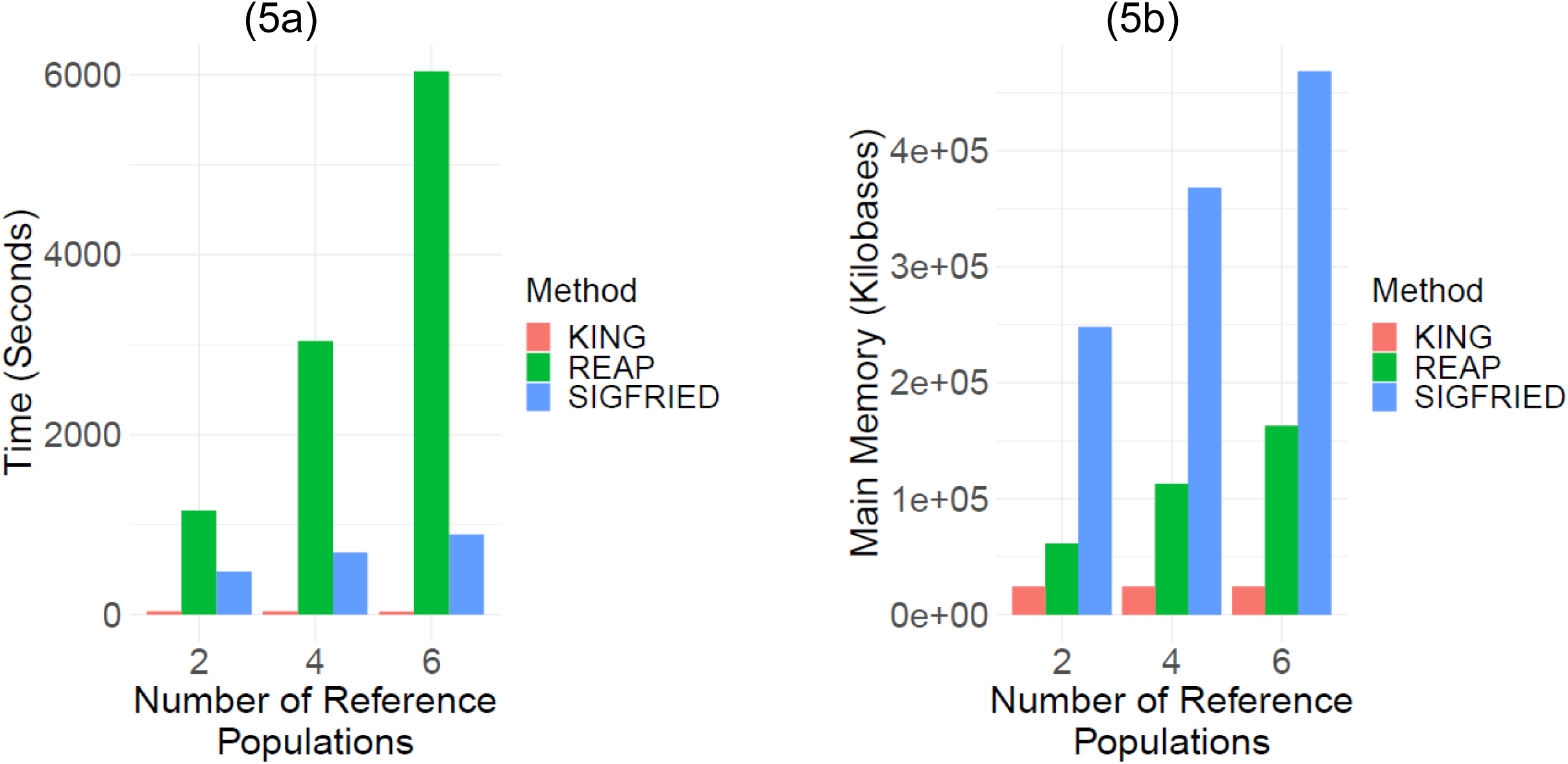
Time and memory requirements of kinship estimation. **(a)** Time requirements (y-axis) by different methods. **(b)** Memory usage (y-axis) by kinship estimation methods.

### Secure Federated Estimation of Kinship Statistics in Two-Site Setting

One of the main advantages of SIGFRIED over previous approaches is enabling privacy-aware kinship estimation in different scenarios due to its modular formulation. We focus on a 2-site collaborative scenario (such as genealogy companies or two institutions working under different regulations) where the sites aim at computing the pairwise kinship statistics among the collective set of individuals in two sites but they cannot share genotype data in plaintext format because of local privacy requirements. We also assume that the sites behave honestly without collusions or malicious data manipulations [62]. This scenario is illustrated in Figure 6a. The two sites have the genotypes matrices *G*^(1)^ and *G*^(2)^ for *S*_1_ and *S*_2_ individuals. The main task is to compute the kinship coefficients between all pairwise comparisons of *S*_1_ and *S*_2_ individuals among the sites. We assume the sites utilize the same reference panels to perform projection-based estimation of admixtures and the individual specific AFs for each individual locally.

**Figure 6.**
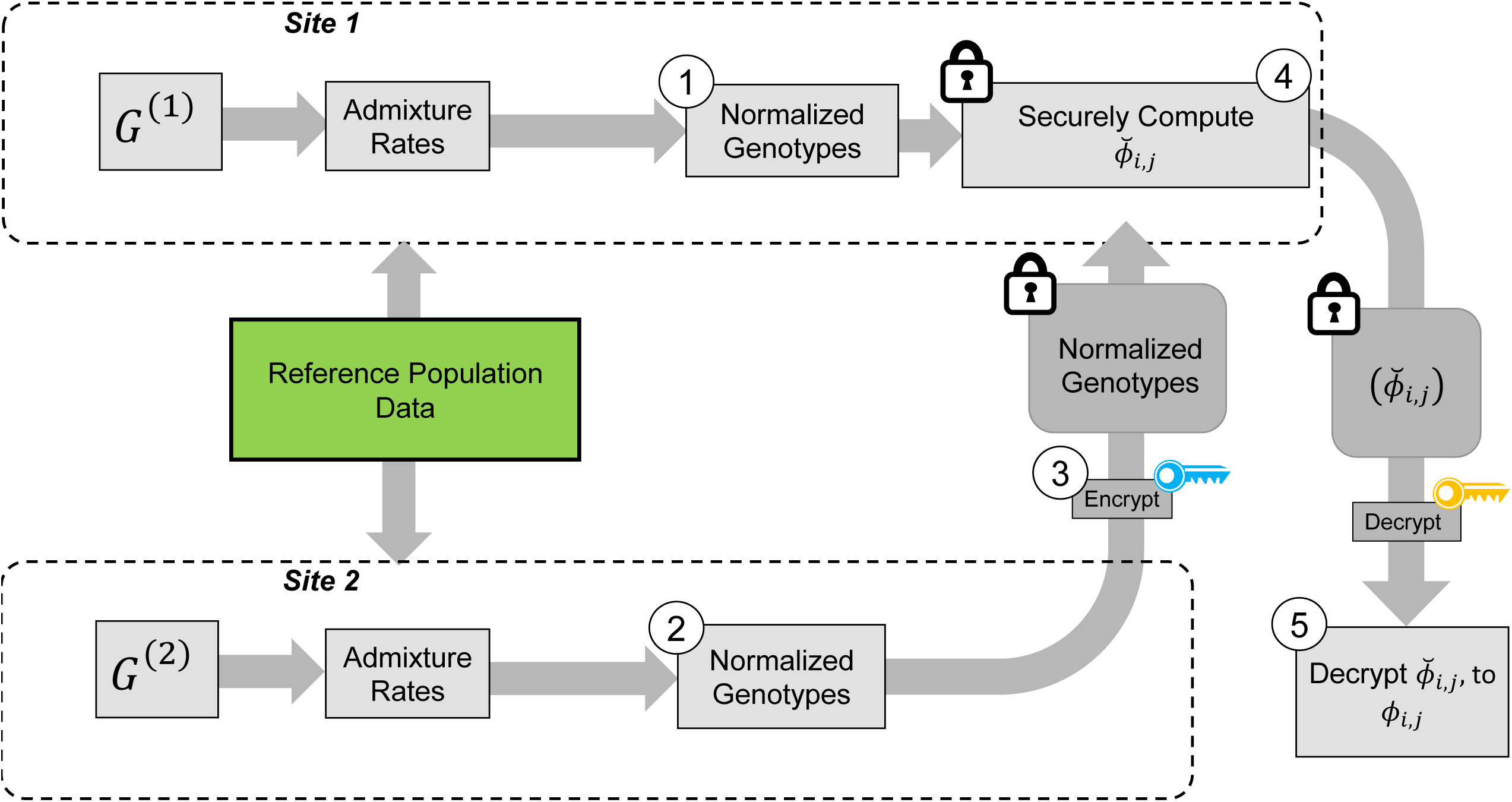
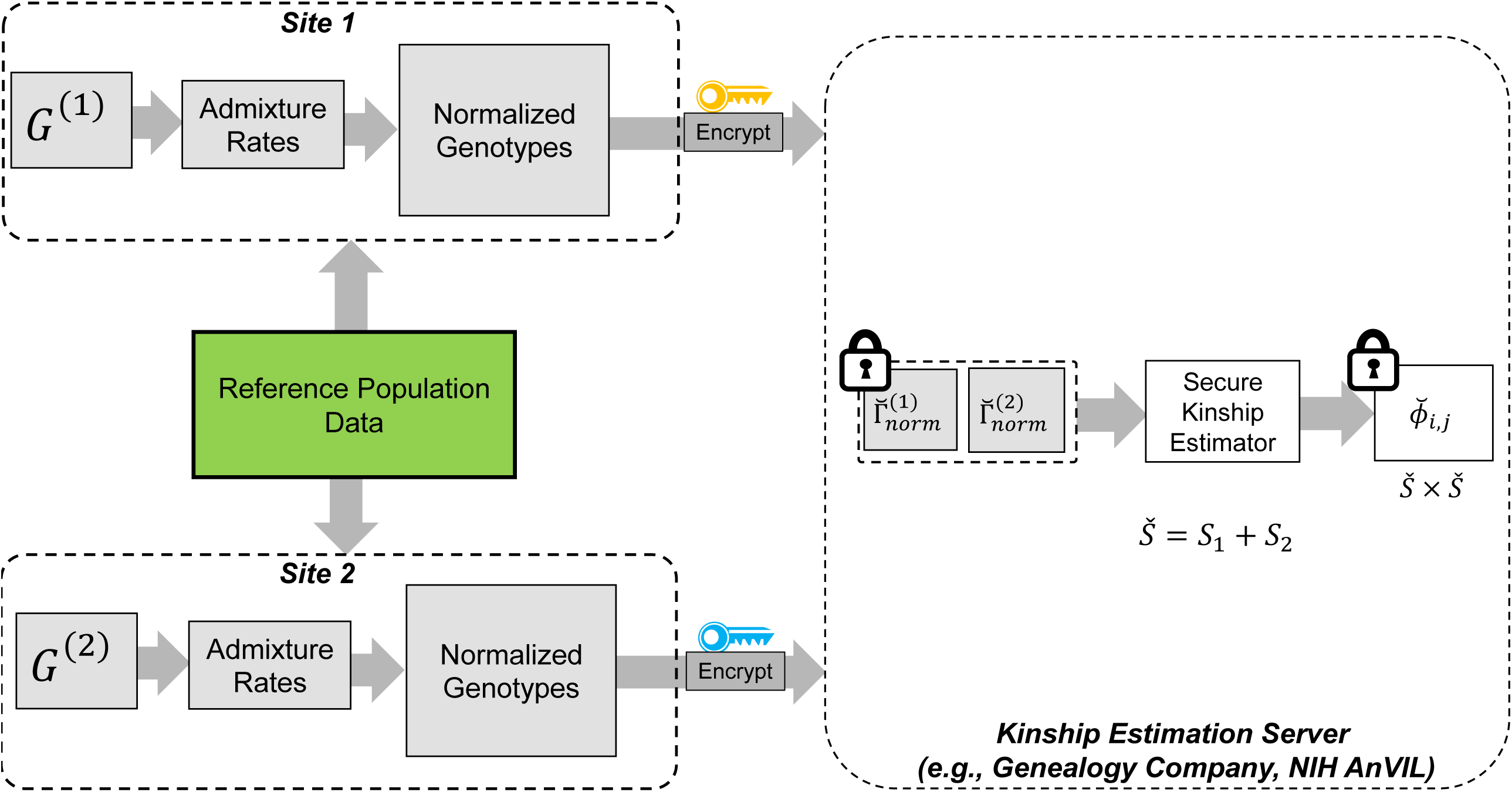
Illustration of secure kinship and IBD-Sharing probability estimation for 2-Site collaboration. **(a)** Site-2 computes the normalized genotypes 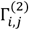 and sends them to Site-1 after encrypting them with the public key. Site-1 also compute the normalized genotype matrix, 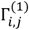. After receiving the encrypted genotype matrix from Site-2, Site-1 securely estimates the encrypted kinship 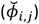 and other statistics. Site-1 sends the encrypted matrices to Site-2, which decrypts the kinship statistics and shares them with Site-2. **(b)** Illustration of a secure kinship estimation with an outsourcing server. The two sites compute normalized genotype matrices. The sites encrypt genotype matrices and send them to the kinship estimation server. The server pools all encrypted data and securely estimates kinship statistics among all samples. The encrypted kinship statistics are then sent to each site, each of which decrypt the kinship statistics.

#### Secure Computation of Correlation-based Kinship Coefficient

We first compute a normalized genotype matrix for each site, Γ^(*a*)^, which denotes the normalized genotype matrix for site *a* by correcting with respect to allele frequencies. *ϕ*_*i,j*_ is computed from the normalized genotype matrices. An important observation is that normalized genotype matrices in each site can be computed locally and do not depend on the other site’s private information. However, this still requires the sites to share the normalized genotype matrices in plaintext format with each other. It is therefore necessary to protect at least one of the matrices by encryption (Fig. 6a). We make use of *homomorphic encryption* to secure the data [61], which enables the processing of the encrypted data without decrypting it. In this setup, both sites compute the normalized matrices and Site-2 homomorphically encrypts and sends its encrypted genotype matrix to Site-1 (or vice versa). We denote the encrypted normalized genotype matrix of Site-2 with 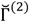. After Site-1 receives encrypted genotypes, it computes the kinship coefficient securely using Г^(1)^ and 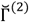. Finally, the computed kinship estimates are sent back to Site-2, which decrypts and shares the kinship coefficient matrix with Site-1.

#### Other Kinship Statistics

Other kinship statistics such as zero-IBD sharing probabilities and distance-based kinship estimator can be securely calculated using an approach similar to as described above under different scenarios.

#### Time and Memory requirements

We implemented a 2-Site kinship estimation using the SEAL library [63]. We used the CKKS encryption scheme with default security settings that satisfy 128-bit security requirements [64]. We used 100 simulated individuals and used 15,000 variants whereby each individual’s genotypes fit into multiple ciphertexts. This proof-of-concept implementation finished the computation kinship coefficients in under 1 minute using less than 4 gigabytes of main memory, which includes encryption, encoding, evaluation, decoding, and decryption time. Overall, we observed that the maximum absolute difference between plaintext and encrypted kinship coefficients is 10^−9^, which practically does not cause differences in analysis of relatedness. The secure computations can be extended to an optimized version of secure federated kinship statistics in multi-site with and without an untrusted outsourcing entity such as a cloud-based server (Fig. 6b). As the data encryption is implemented in our protocol, even untrusted entities can be used in federated kinship estimation for making use of large cloud-based scaling for improved performance [65,66].

## Discussion

Kinship and related statistics are essential in many genetic studies and they are sensitive for individual and group-level privacy. Here, we presented SIGFRIED, an efficient, accurate, and secure method that utilizes projection on existing reference panels. SIGFRIED balances accuracy and efficiency to ensure that the final algorithm is efficiently implemented with secure primitives. While projection on existing population panels has been utilized previously by other methods, SIGFRIED utilizes projection to circumvent computations that are otherwise hard to implement in the secure domain. From this perspective, we view SIGFRIED as a private-by-design methodology wherein the privacy considerations are balanced against efficiency and accuracy. Projection does not explicitly require reference panel genotypes. Since the reference genotypes are not explicitly shared, this creates minimal risk for reference panels under restricted access (i.e. TOPMed [67]).

While we presented a specific privacy-preserving scenario for a 2-site federated estimation of kinship, the implementation and the scenarios can be differently set up to expand to more than 2 sites and also for utilizing an outsourcing service for kinship estimation. The outsourcing can be performed by an untrusted entity because sensitive data is encrypted and cannot be used to infer any information by an unauthorized party. When deployed on a highly scalable but untrusted environment such as AWS or Google Cloud, the performance can be tuned as desired. Also, SIGFRIED implements kinship estimation using modular steps and decomposable functions. This is beneficial for optimizing privacy-vs-performance in different scenarios. The modularity is important because new protocols can choose to encrypt only certain parts of the intermediate statistics to ensure that performance is optimized and security requirements are met according to local regulations and patient or participant consent. For instance, the individual-specific allele are highly aggregated functions of genotypes and can be deemed safe to share in plaintext form.

## Methods

### Variant Selection and Simulations

For simulations, we filtered the variants in The 1000 Genomes Project by first selecting the variants with minor allele frequencies greater than 5% on the autosomal chromosomes 1 through 22. Next, we divided the samples with respect to their assigned populations and used these as population specific reference panels. Simulations are performed by sampling variants for each individual with respect to allele frequencies and the relatedness.

### Projection-based Estimation of Kinship Statistics

Figure 1 summarizes the kinship estimation approach by SIGFRIED. Kinship estimation takes a query genotype matrix, *G*_*N*×*S*_, that contains the genotypes of *N* variants for *S* individuals. The output is *S* × *S* matrix of kinship related statistics. The kinship statistics are calculated and implemented by the formulations that are presented in the Results Section using correlation and distance-based statistics.

## Source Code and Data Availability

Source code and Datasets will be made available upon publication of this manuscript.

## Notes

### Competing Interest Statement

The authors have declared no competing interest.

